# PCTA, A PAN-CANCER CELL LINE TRANSCRIPTOME ATLAS

**DOI:** 10.1101/2024.01.10.575087

**Authors:** Siyuan Cheng, Lin Li, Xiuping Yu

**Affiliations:** Department of Biochemistry & Molecular Biology, LSU Health Shreveport; Feist Weiller Cancer Center, LSU Health Shreveport; Department of Urology, LSU Health Shreveport

## Abstract

**Background:** A substantial volume of RNA sequencing data were generated from cancer cell lines. However, it requires specific bioinformatics skills to compare gene expression levels across cell lines. This has hindered non-bioinformaticians from fully utilizing these valuable datasets in their research. To bridge this gap, we established a curated Pan-cancer Cell Line Transcriptome Atlas (PCTA) dataset. This resource aims to provide a user-friendly platform, allowing researchers without extensive bioinformatics expertise to access and leverage the wealth of information within the dataset for their studies. Importantly, PCTA stands out by offering sufficient sample numbers per cell line in comparison to other pan-cancer datasets.

**Methods:** Cell lines’ meta data and RNA sequencing data were retrieved from the Cancer Cell Line Encyclopedia (CCLE), SRA and ARCHS4 databases. Utilizing the programming language R, we conducted data retrieval, normalization, and visualization. Only expression data for protein-coding genes and long-non-coding RNAs (LncRNAs) were considered in this study, streamlining the focus to enhance the precision and relevance of the analysis.

**Results:** The resulting PCTA dataset encompasses the expression matrix of 24,965 genes, featuring data from 84,385 samples derived from 5,677 studies. This comprehensive compilation spans 535 cell lines, representing a spectrum of 114 cancer types originating from 30 diverse tissue types. On UMAP plots, cell lines originating from the same type of tissue tend to cluster together, illustrating the dataset’s ability to capture biological relationships. To unravel molecular signatures, marker genes were identified for each cancer type. Additionally, an interactive and user-friendly web application (https://pcatools.shinyapps.io/PCTA_app/) was developed for researchers to explore the PCTA dataset. This platform allows users to examine the expression pattern of their genes of interest across a diverse array of samples. Data are visualized as violin-, box-, and point-plots, enhancing the interpretability of the findings.

**Conclusion:** The PCTA stands as a comprehensive resource, offering insights into gene expression patterns across diverse cancer cell lines and providing a valuable tool to explore molecular signatures and potential therapeutic targets in cancer research.

## INTRODUCTION

Cancer accounts for 1 in 6 deaths globally^1,2^. Urgent efforts are underway to identify effective anti-cancer treatments. Established cancer cell lines are pivotal in cancer research, serving as the most widely used models. These cell lines are relatively more cost-efficient and user-friendly compared to alternatives such as mouse models or patient-derived xenografts (PDXs). Notably, most cancer cell lines exhibit greater genomic and transcriptomic homogeneity than human primary tumors, making them valuable tools for unraveling molecular mechanisms following genetic perturbations or drug treatments. Despite their crucial role, cancer cell lines remain under-characterized, especially in the context of comparing gene expression across diverse cell lines. Existing pan-cancer cell line studies, such as the Cancer Cell Line Encyclopedia (CCLE)^3^ and NCI-60^4^, provide transcriptome profiles but with a limited number of samples.

We have previously launched a highly acclaimed web application, the <u>C</u>ombined <u>T</u>ranscriptome dataset of human <u>P</u>rostate cancer <u>C</u>ell lines (CTPC)^5^, which has garnered widespread recognition and is consistently accessed by users worldwide. The positive reception of this tool underscores the research community’s keen interest in a user-friendly platform for visualizing gene expression across diverse samples. Drawing inspiration from the success of CTPC, we introduce the <u>P</u>an-cancer <u>C</u>ell Line <u>T</u>ranscriptome <u>A</u>tlas (PCTA). This resource is designed to provide researchers with valuable insights into gene expression patterns across commonly used cancer cell lines, spanning various cancer types. In line with the methodology established in the CTPC study, we stress the importance of an ample number of samples, enabling the median gene expression value to accurately reflect the expression status across all samples from the same cell line. Furthermore, outlier values can shed light on treatment-induced gene expression perturbations. Our data collection pipeline is designed to facilitate periodic updates, ensuring the inclusion of the latest information, and we envision the potential expansion of the collection to encompass multi-omics data.

## METHODS

### 1. Data collection and filtering

The list of commonly used cancer cell lines and their metadata including cancer types and tissue origins were obtained from CCLE^3^. Subsequently, the names of these cell lines were used as keywords to extract relevant samples’ metadata from the Sequence Read Archive (SRA) database. This retrieval process was executed through a combined function of “esearch” and “efetch,” utilizing the “Entrez Direct” toolkit via Unix terminal commands^6^.

The resulting metadata file, encompassing approximately one million samples, was imported into the R environment for the purpose of selecting the RNAseq samples that were generated using the respective cell lines. The samples were selected based on the following specific attributes: the sample’s taxonomy ID as “9606” (human), library strategy as “RNA-seq”, library source as “Transcriptomic”, and the sample’s accession name contains “GSM” which indicate retrievable metadata from GEO database. The corresponding keyword used for fetching samples could be present in either the sample name or sample description. Following this filtering process, approximately 160K samples remained. Since the “efetch” function did not retrieve all experimental metadata for each sample, the “getGEO” function from “GEOquery” R package^7^ were used to download comprehensive metadata using the filtered GEO accession numbers.

The raw read counts at the transcript level were obtained from ARCHS4^8^ (Ensembl 107 version) using customized R code. Subsequently, the count value of each transcript was transformed into Transcripts Per Million (TPM) using the “convertCounts” function from “DGEobj.utils” R package^9^. The transcript length information was acquired from Ensembl GRCh38. After that, the transcript-level TPM values were aggregated to generate gene-level TPM values by summing the values of transcripts belonging to the same gene.

### 2. Data processing, visualization and the construction of PCTA web application

Customized R codes based on “Seurat”^10^ and “SCpubr”^11^ packages were used to perform dimensional reduction, calculation of marker genes and data visualization. Marker genes specific to various cancer types were determined using the “wilcox” method with gene counts. Additionally, average TPM values were calculated based on gene TPM values for each cancer type. To filter marker genes, two criteria were applied: a log2 TPM mean higher than 5 and a log2 TPM standard deviation lower than 3. The R “ggplot2” package^12^ was used to generate violin plot, box plot and jitter plot (a variant of point plot) for effective data representation.

The ShinyApp platform was used to host the custom R codes for PCTA web application. To facilitate data normalization, TPM values of each gene were transformed to log2 TPM +1 value. Only expression data for protein-coding genes and LncRNAs were included in the PCTA website to optimize processing efficiency.

## RESULTS

### Overview of the Pan-cancer Cell Line Transcriptome Atlas (PCTA) Dataset

Following the workflow depicted in Figure 1A, the PCTA dataset was established, encapsulating gene expression matrices for 84,385 samples across 24,965 genes. To capture the gene expression landscape comprehensively, two distinct formats were generated and used, raw counts and Transcripts Per Million (TPM). These samples, derived from 5,677 studies, spanned 535 cancer cell lines, representing 114 cancer types originating from 30 distinct tissue types (Figure 1B). The annotated list of 535 cell lines includes details regarding their anatomic origin and cancer type, with cancer type abbreviations adapted from CCLE and available on OncoTree website^13^ (https://oncotree.mskcc.org/#/home).

**Figure 1.**
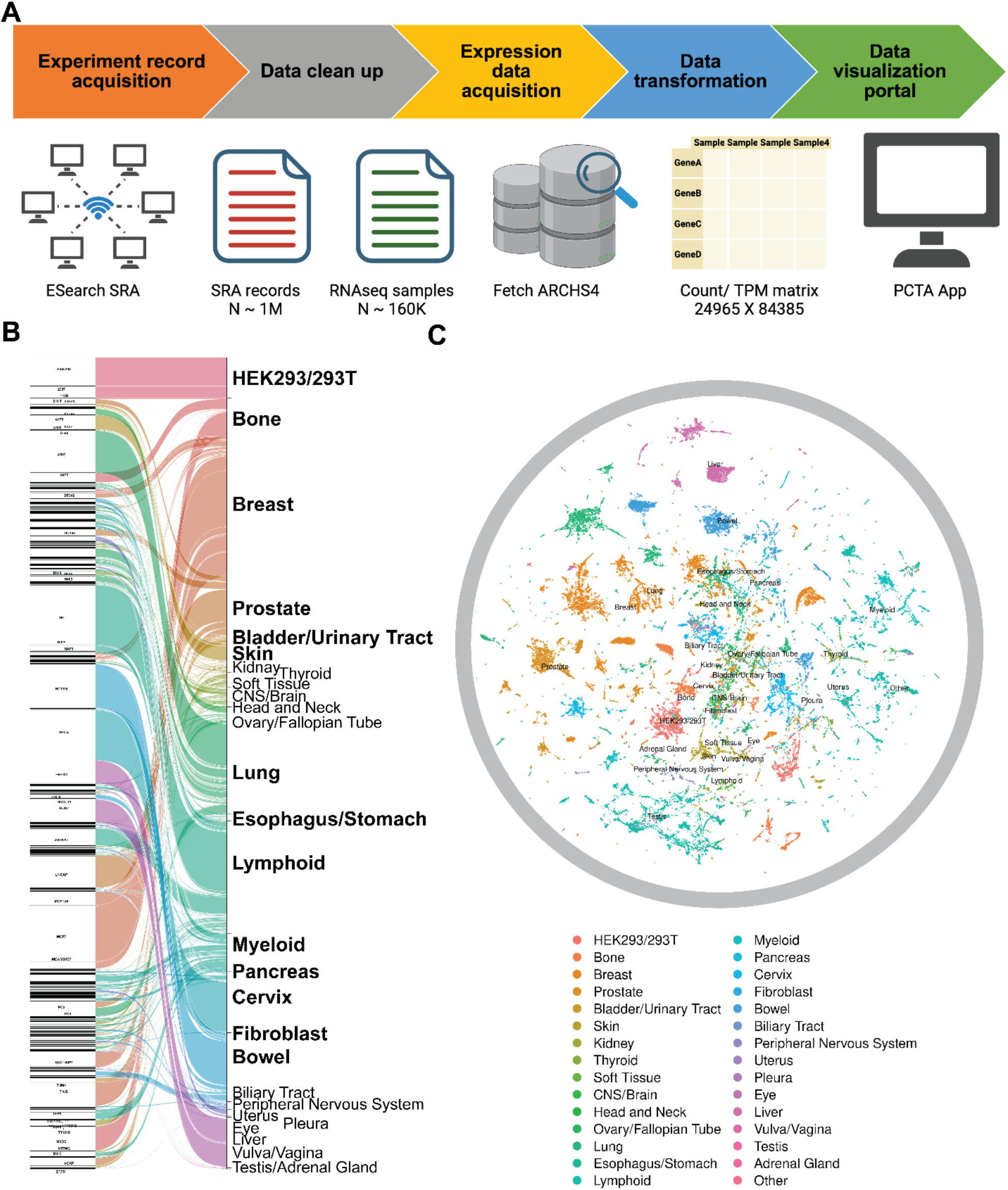
Establish of PCTA database. The carton charts was used to demonstrate the pipeline to build PCTA database (A). The sanky plot showed the relationship between PCTA cancer cell lines and their anatomic origins (B). The UMAP showed the relatively clustered location of samples from the same cancer origin (C).

### Visualizing Transcriptomic Patterns

UMAP (uniform manifold approximation and projection) plots were employed to visually convey the transcriptomic profiles of each PCTA sample. These plots unveiled close clustering of samples originating from the same cancer origins (Figure 1C), cancer types (sFigure 1A), and individual cell lines (sFigure 1B), underscoring the biological coherence within the dataset.

### Identification of Marker Genes

Marker genes specific to each cancer type were discerned, with the top-ranking ones, based on log2 fold change, presented in Table 2. The complete list is available in sTable 1, providing valuable insights into the distinctive genetic signatures characterizing different cancer types.

**Table 1.**
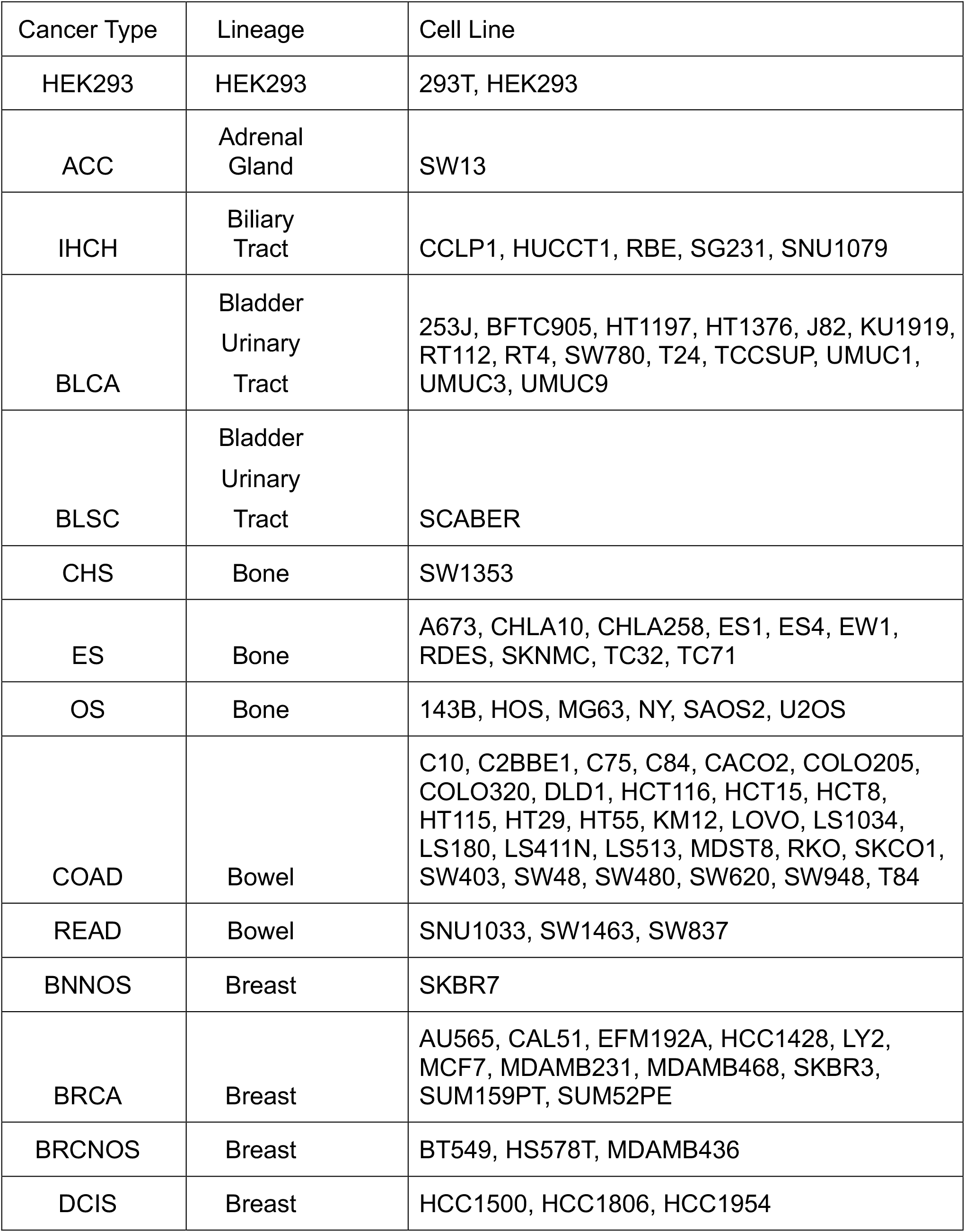
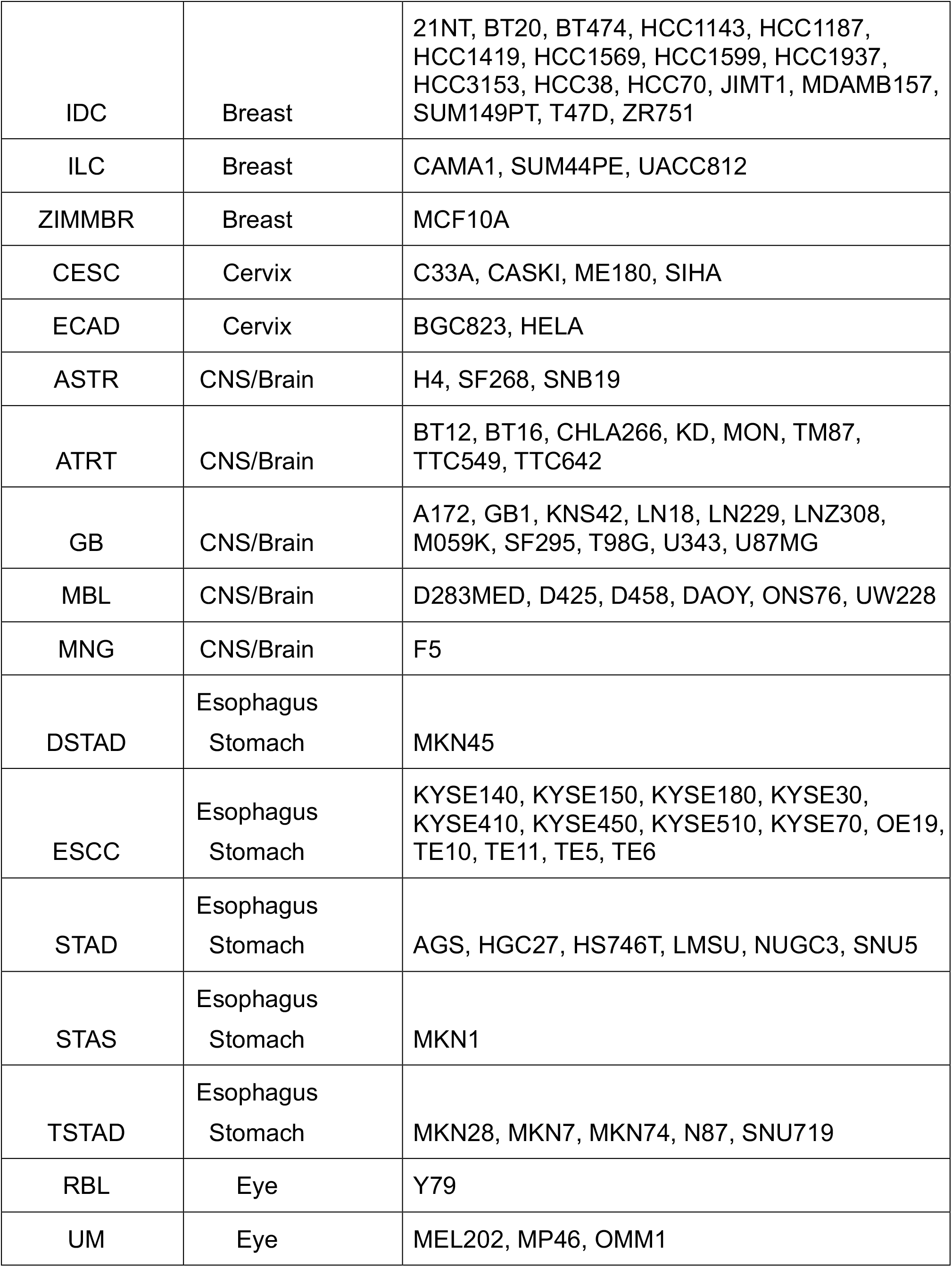
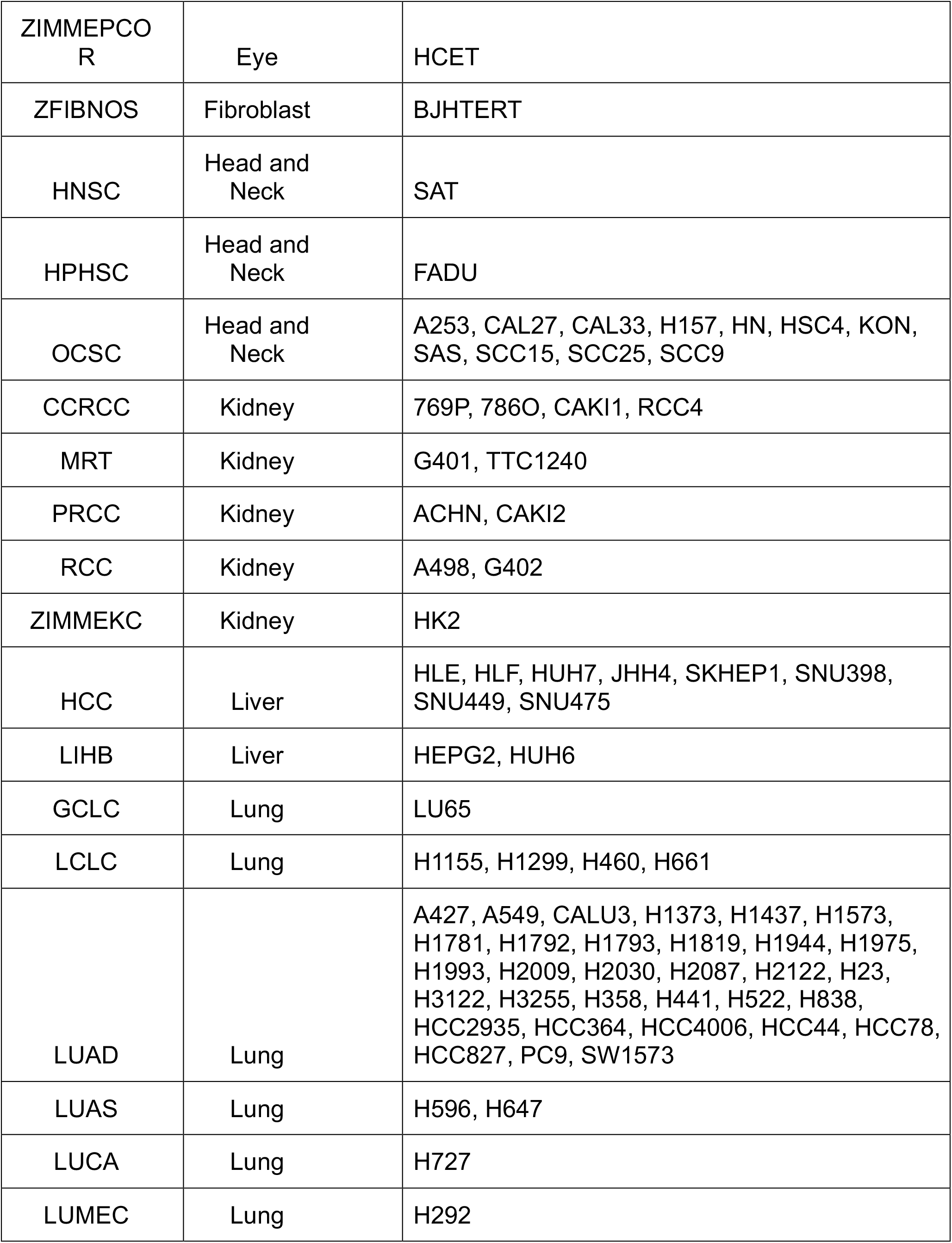
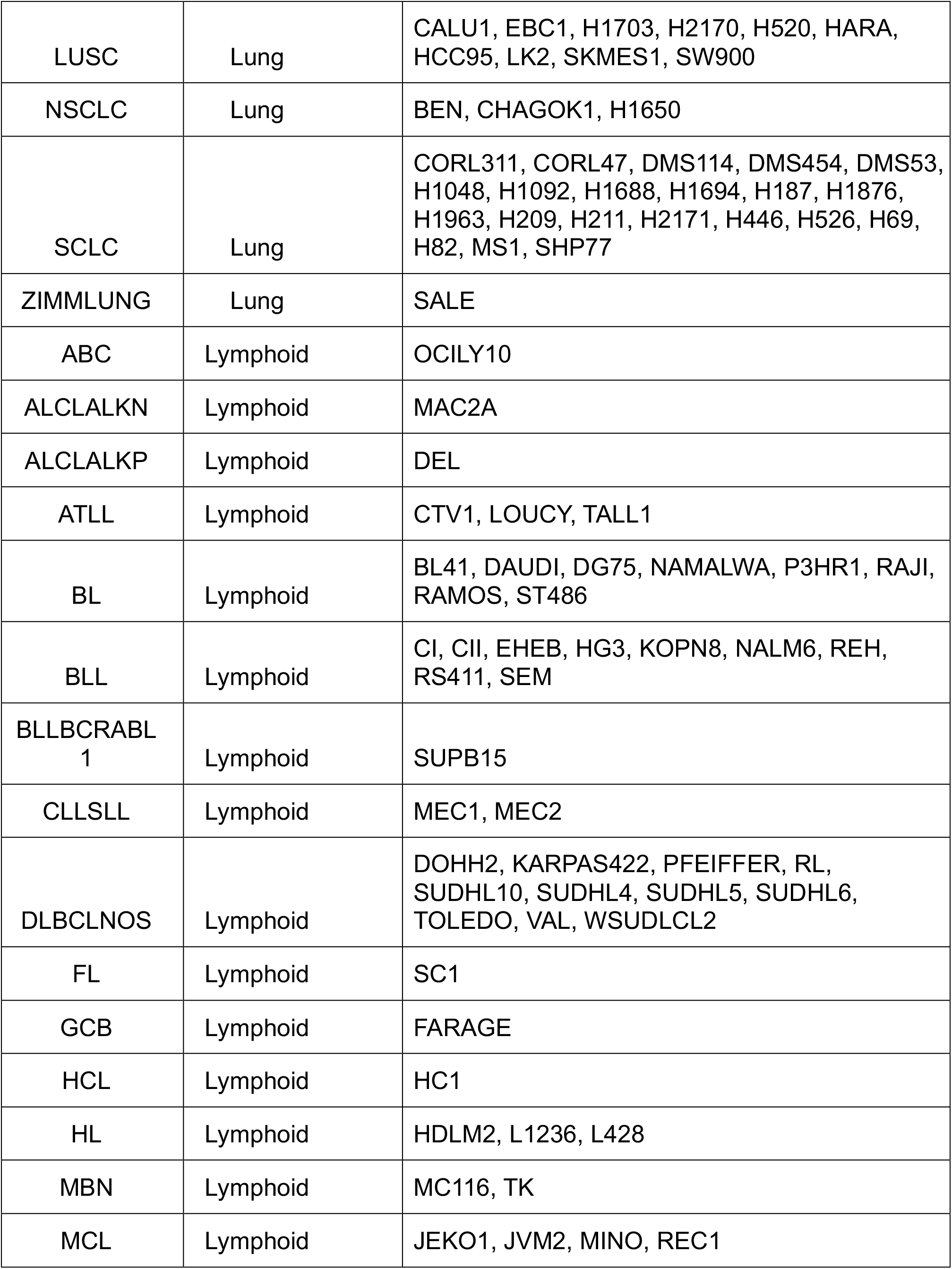
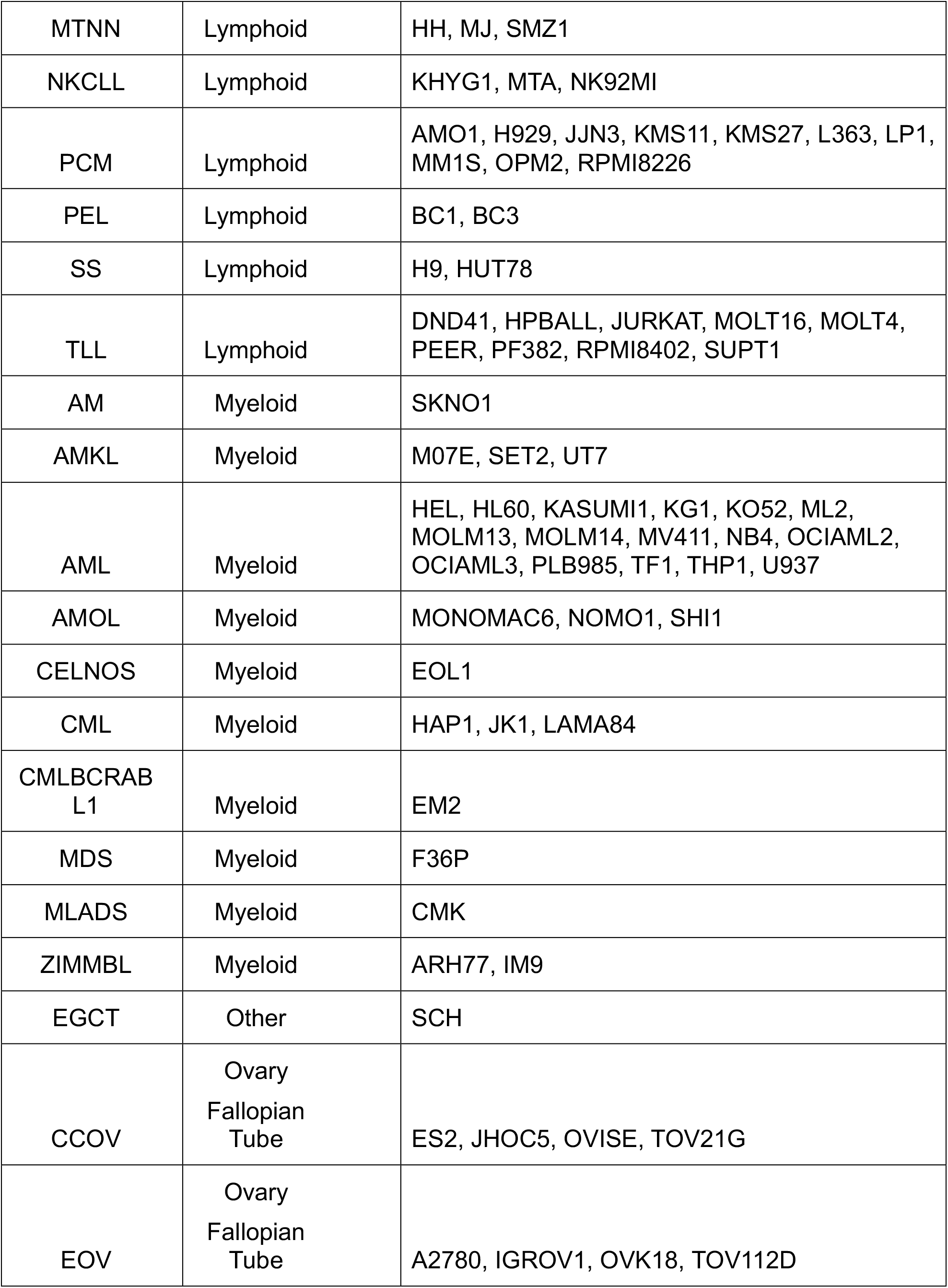
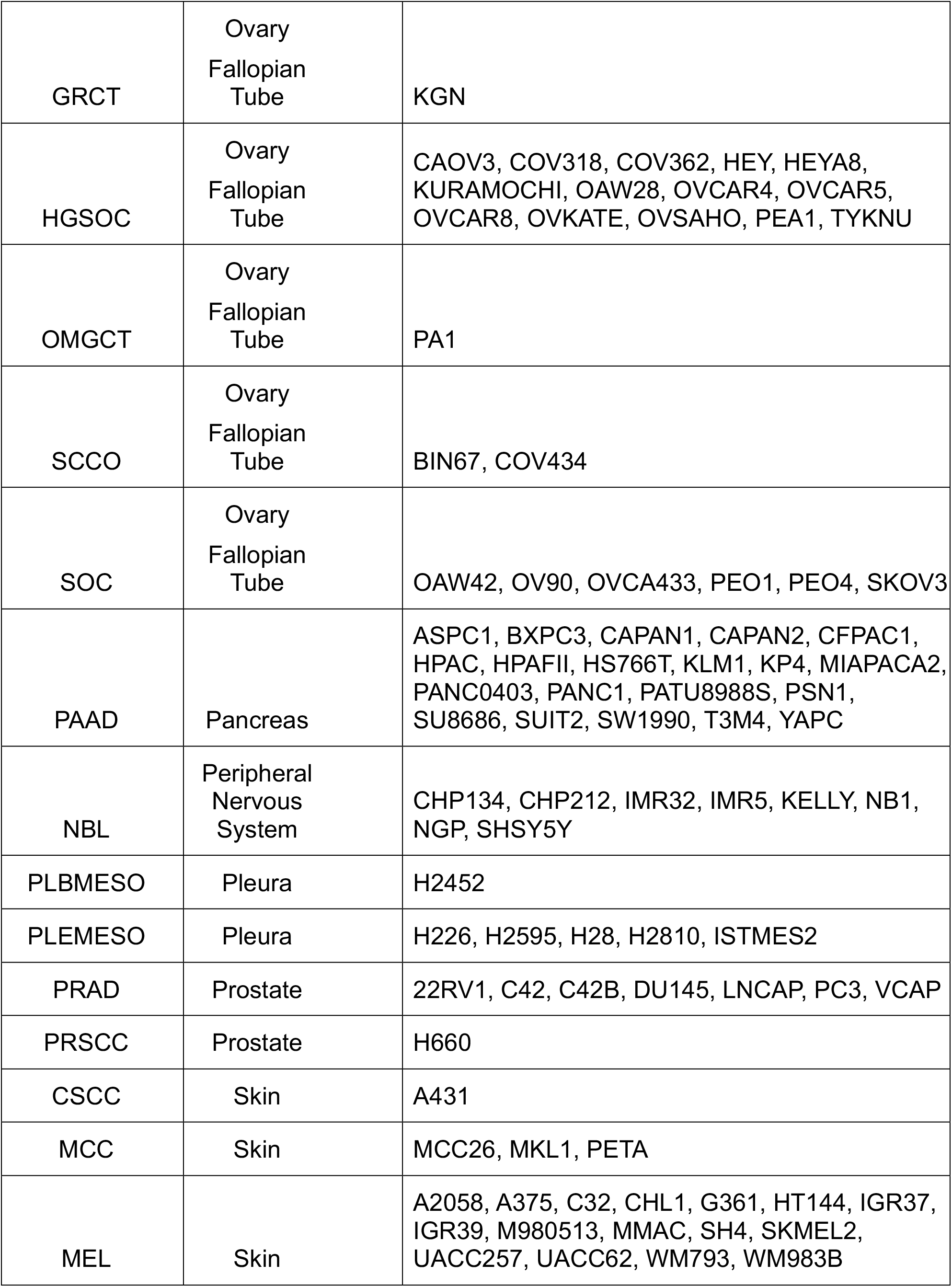
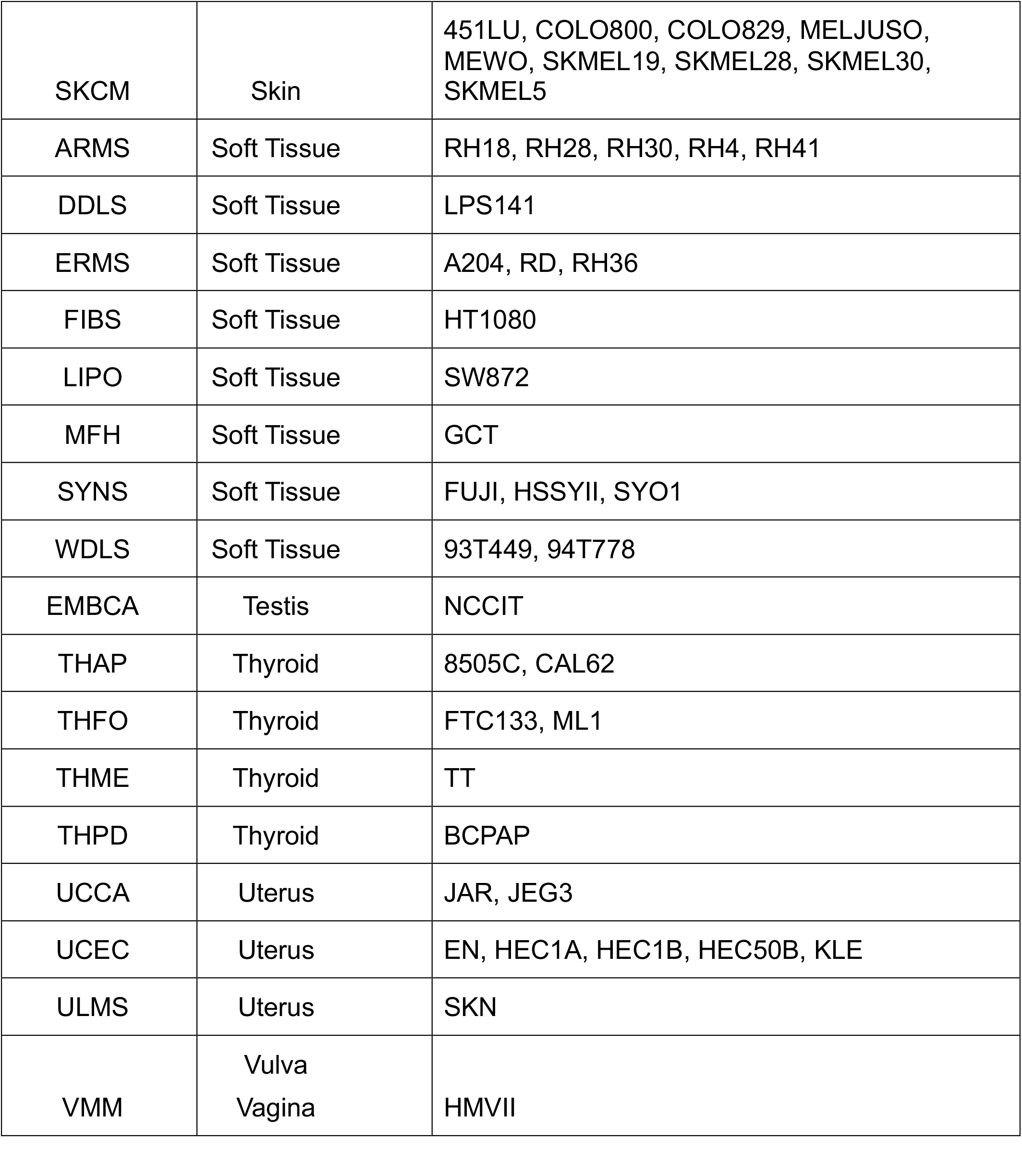
List of PCTA cell lines.

**Table 2.**
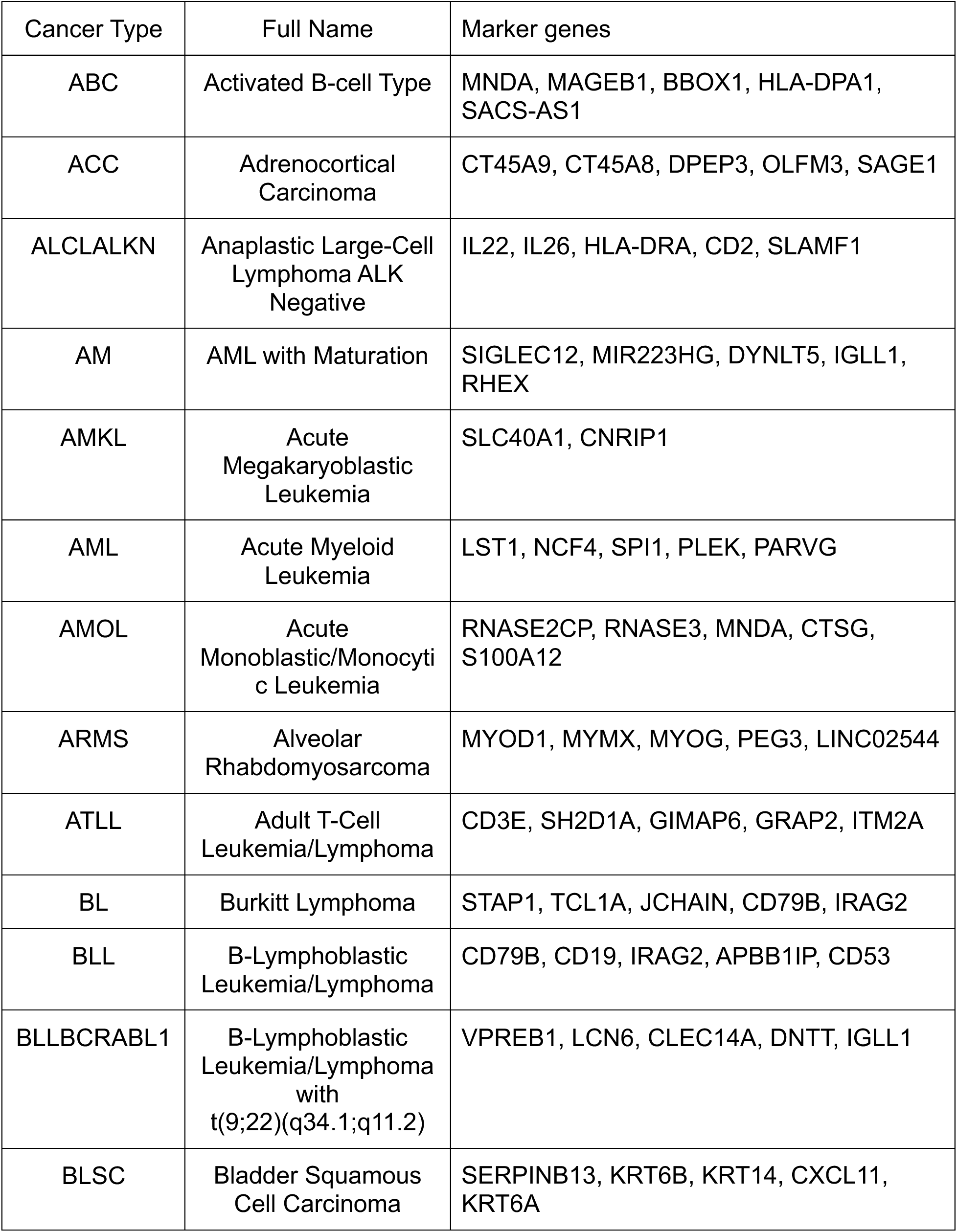
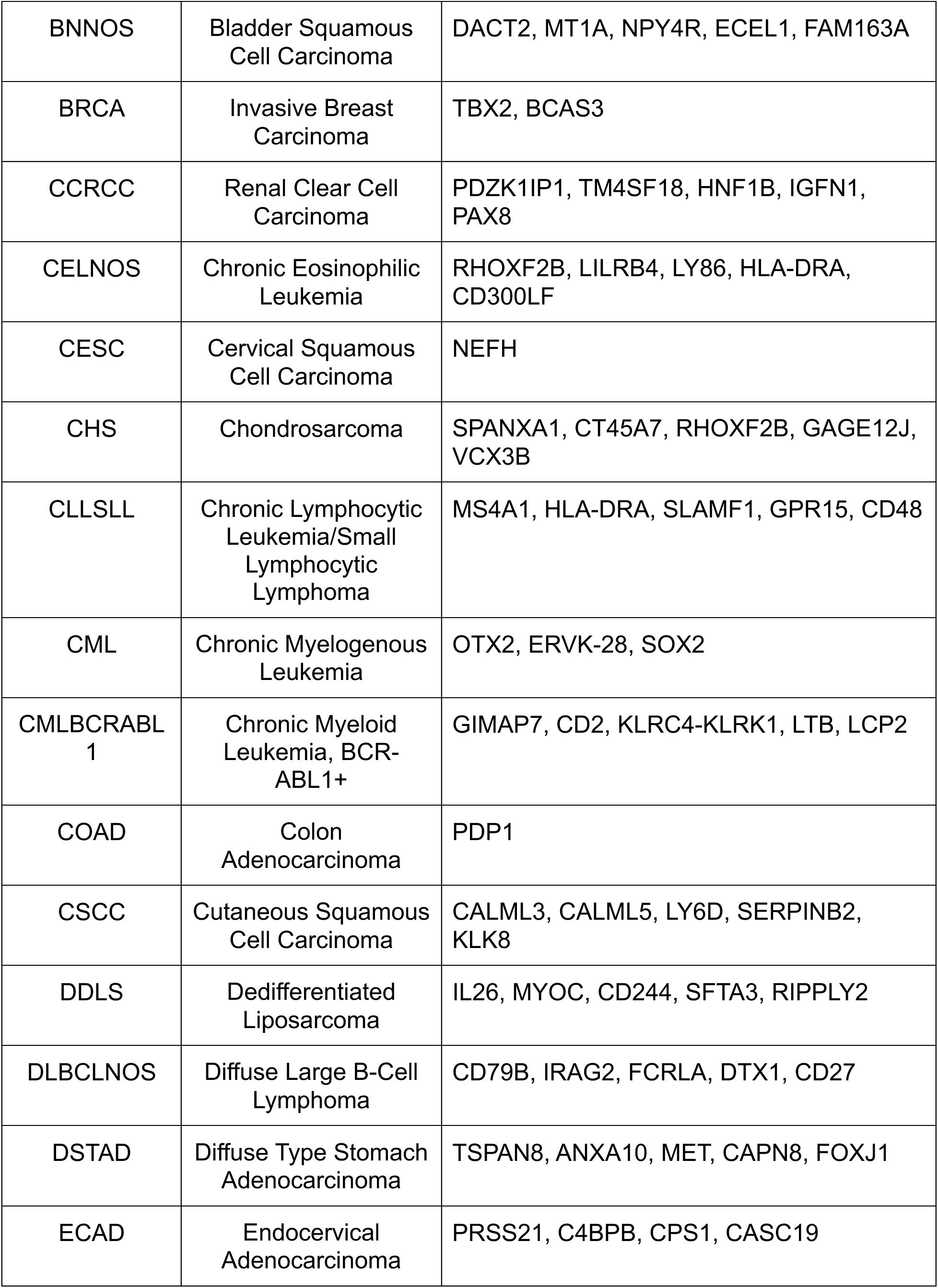
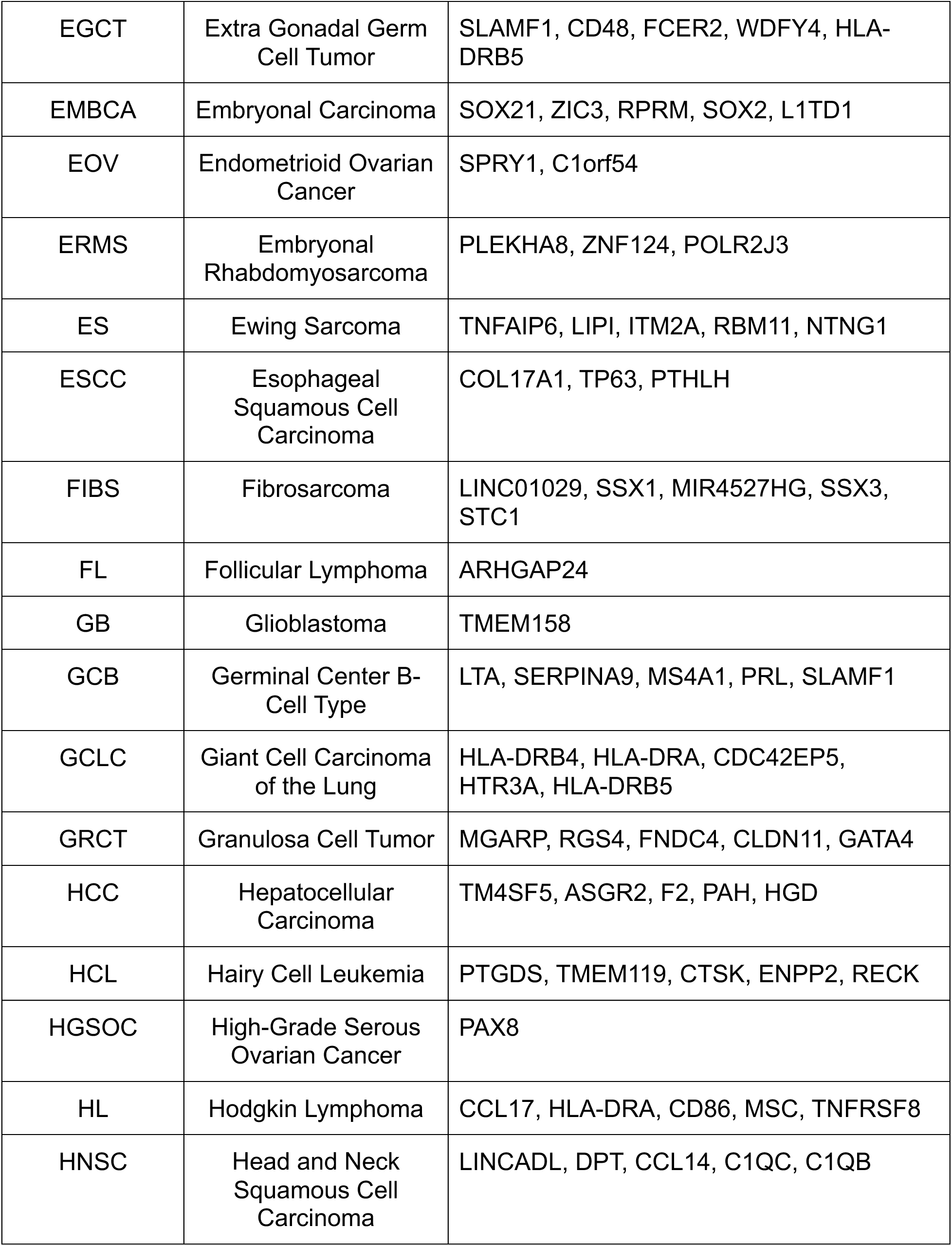
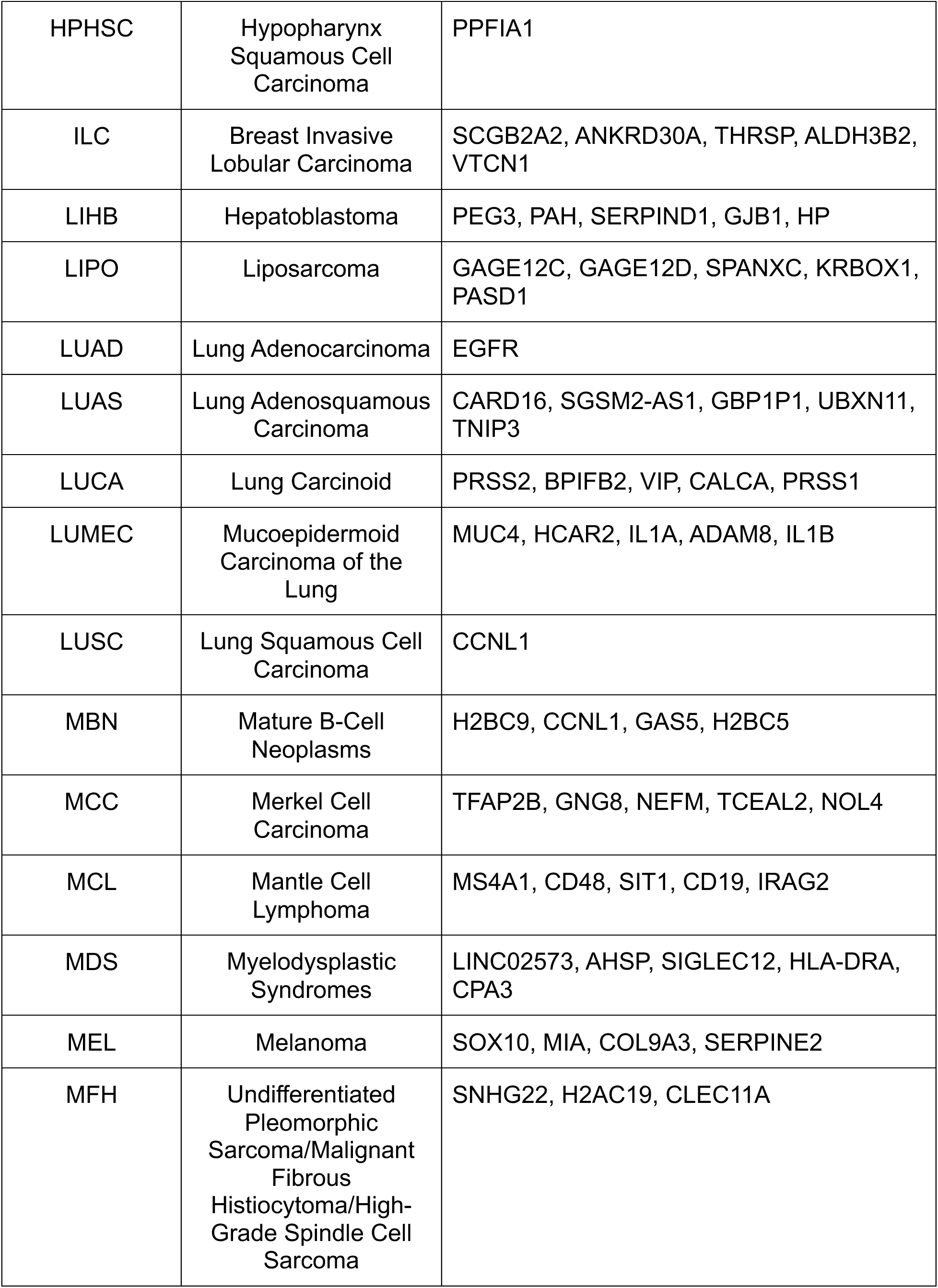
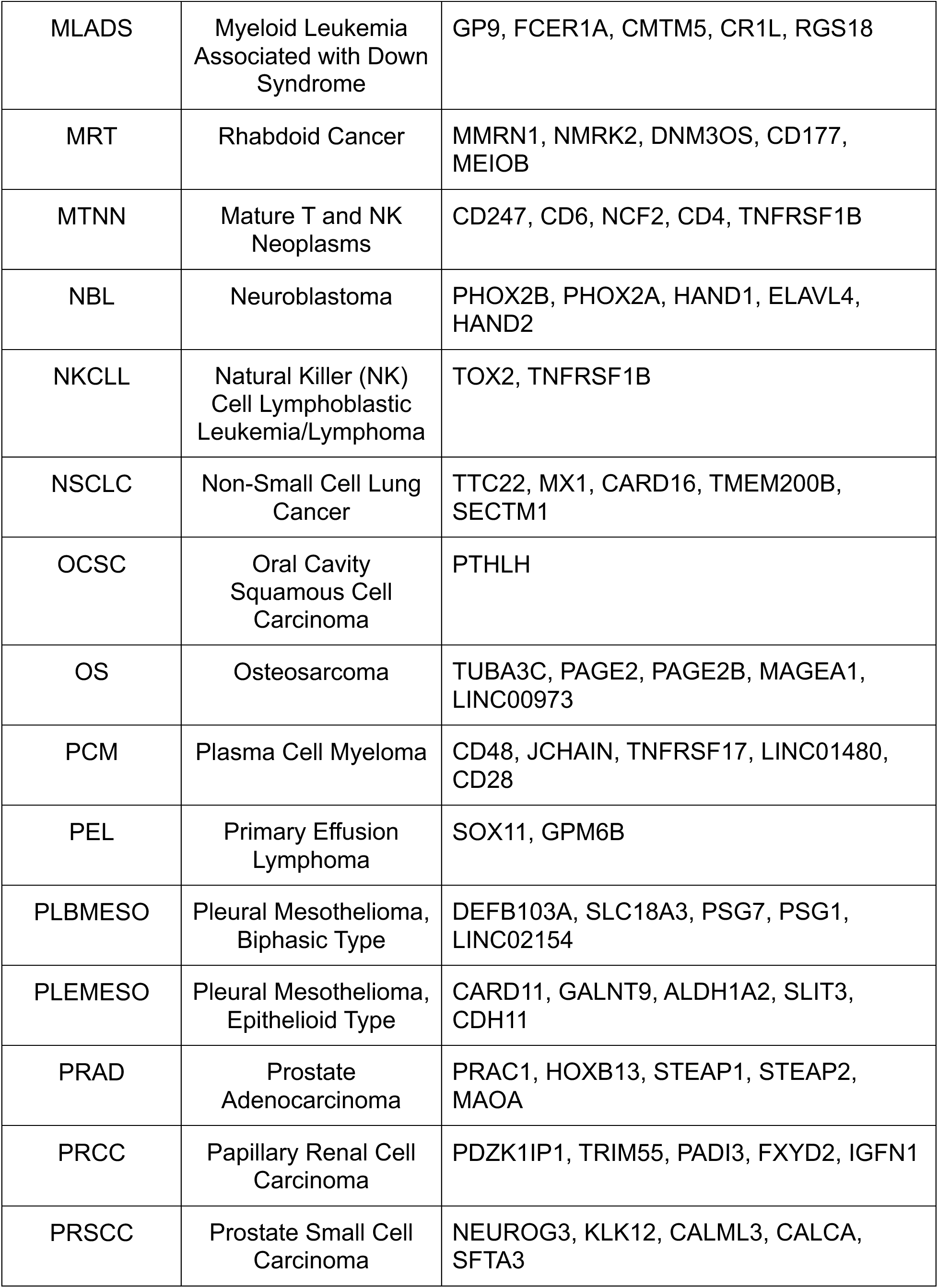
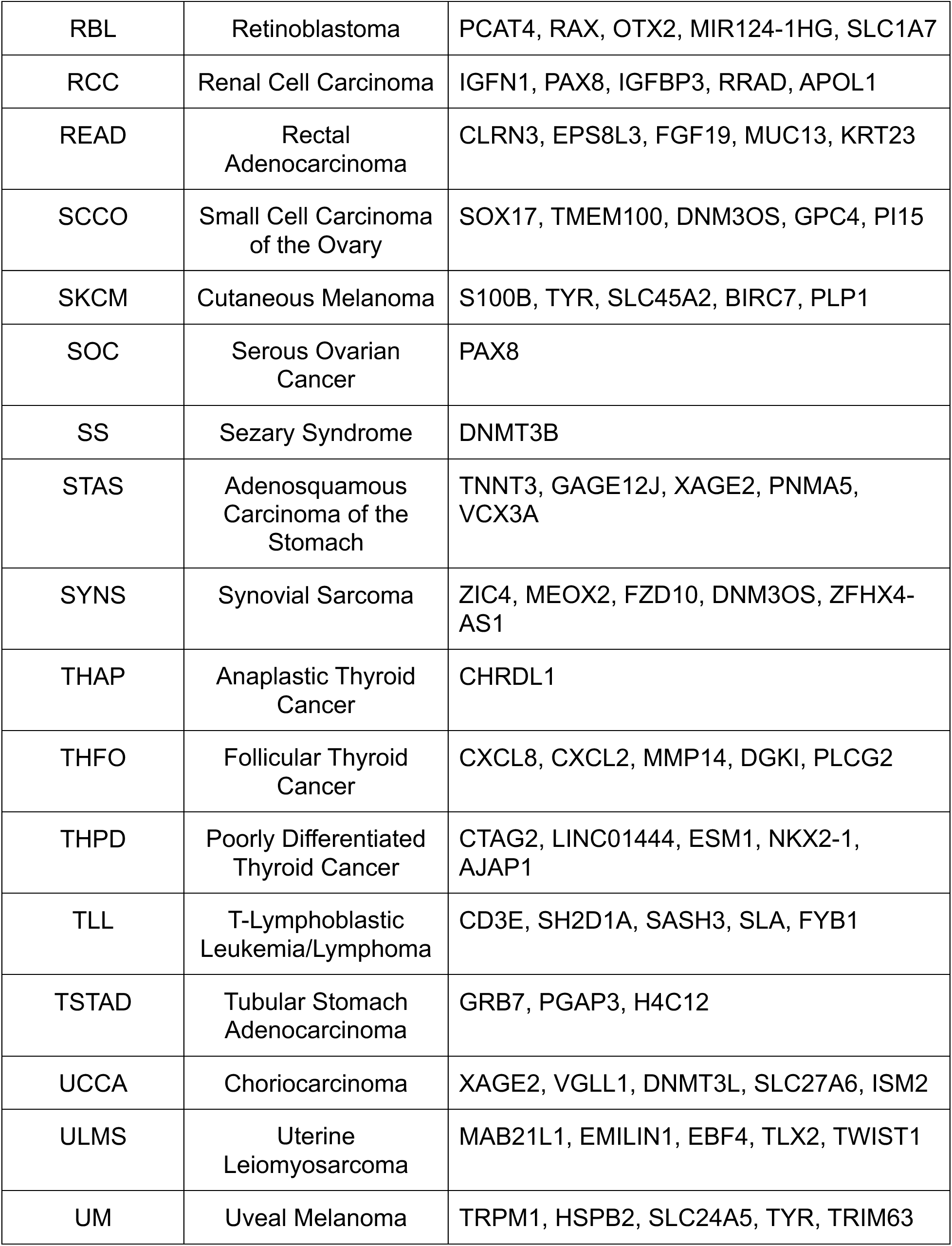
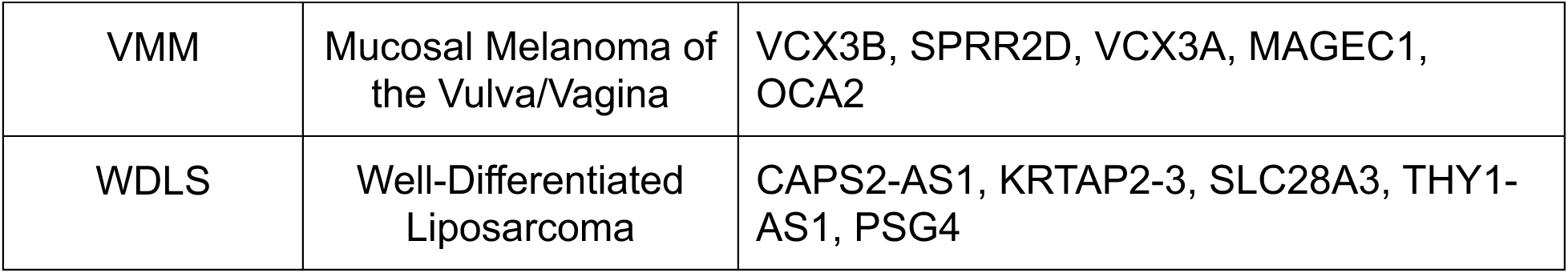
Cancer type marker genes.

### Facilitating Public Access with the PCTA Website

We developed the PCTA web application https://pcatools.shinyapps.io/PCTA_app/ to provide users with access to gene expression data across diverse cancer origins and cell lines. This interactive platform enables users to input the official gene symbol of interest and explore quantification plots. Each gene’s expression is vividly portrayed through a violin plot, offering a comprehensive overview of its distribution across all samples within the PCTA collection, categorized by cancer origin.

Furthermore, the PCTA website empowers users to focus specifically on anatomic origins by selecting corresponding tags. In this customized view, gene expression is visualized through a combination of boxplots and point plots, with samples grouped by cell line names. The interactive nature of this plot allows users to seamlessly zoom in/out and pinpoint specific sample dots for an in-depth exploration of associated metadata, enhancing the precision and depth of user engagement and facilitating a more tailored and insightful analysis of gene expression patterns within specific anatomic contexts.

While the median value of each gene provides a general overview of its expression across cell lines, delving into outlier samples unveils valuable insights for researchers investigating the upstream regulation of a gene of interest. The examination of outlier samples enables researchers to scrutinize treatment conditions associated with significant up- or down-regulation of gene expression. This analytical capability is made possible by the interactive features embedded in the PCTA plots, offering a powerful tool for researchers to navigate and interpret treatment-induced variations in gene expression within the rich context of the Pan-cancer Cell Line Transcriptome Atlas.

## DISCUSSION

Cancer cell lines are pivotal tools for cancer research. PCTA, by furnishing reliable gene expression data across commonly used cell lines, equips researchers with the means to strategically choose cell lines based on the expression patterns of genes relevant to their studies. For instance, leveraging the insights gained from PCTA, researchers can select cell lines for gene manipulation experiments. This may involve overexpressing genes in cell lines where the target genes are expressed at low levels or knocking down genes in cell lines that exhibit high expression levels of the specific genes of interest. This marks a significant advancement in experimental design within the field of cancer research.

In addition to their utility in experimental manipulation, cancer cell lines offer advantages over human specimens, notably in terms of gene expression uniformity. This uniformity arises from the absence of non-cancerous cell populations, rendering cancer cell lines particularly well-suited for discerning markers specific to certain cancer types and potential treatment targets.

The homogeneity in gene expression within cancer cell lines enhances the precision of marker identification, providing a clearer signal amidst the molecular noise. Our findings, particularly in the context of prostate adenocarcinoma (PRAD), highlight several marker genes, such as STEP1, STEP2, and HOXB13, which have been extensively studied and documented. These well-established markers serve as valuable reference points, validating the reliability of the PCTA dataset.

Furthermore, our exploration has revealed novel marker genes, such as PRAC1, which exhibits a robust and specific expression pattern in PRAD and warrants in-depth exploration.

Besides its current version, the PCTA is accompanied by a meticulously crafted pipeline that extends its value to the broader research field. Our pipeline is strategically designed not only to streamline data collection but also to automate data filtering, thereby minimizing human effort and ensuring the stability and frequency of updates. In the ever-evolving landscape of cancer research, where hundreds of samples are deposited into databases like GEO daily, the automated update feature of the PCTA pipeline is invaluable. This capability positions the PCTA to expand dynamically by incorporating new samples, maintaining its relevance and comprehensiveness in capturing the diverse molecular profiles of cancer cell lines.

The versatility of our pipeline extends beyond transcriptomics. It harbors the built-in potential to seamlessly enlarge the collection to include multi-omics data such as ChIP-seq, GWAS and proteomics.

A limitation of the current pipeline stems from its trade-off between computational efficiency and sample breadth. In an effort to optimize computing time and expedite result delivery to users, the PCTA acquires raw read counts from ARCHS4^8^, bypassing the alignment process using fastq files sourced from the SRA database. While this approach ensures promptness, it imposes a constraint on the number of samples, restricted to those available within the ARCHS4 dataset. To address this limitation and enhance the inclusivity of the PCTA, future iterations of the pipeline may embark on innovative strategies, such as performing alignments using raw sequence data, artificial intelligence based large language models (LLMs) techniques, for meta data cleanup and the integration of a more extensive spectrum of samples.

In conclusion, the current PCTA emerges as a valuable tool for cancer research, offering a robust quantification of gene expression data across a diverse spectrum of cancer types. With a focus on high sample numbers, PCTA provides an expansive resource for researchers seeking comprehensive insights into the molecular landscapes of various cancers. The accessibility of the PCTA is further underscored by the user-friendly PCTA website, which serves as a public portal. This platform empowers researchers to effortlessly download and visualize gene expression data, requiring only the input of the gene symbol. The simplicity and accessibility of the PCTA website aim to facilitate and accelerate research endeavors, ensuring that valuable insights from the extensive dataset are readily available to the broader scientific community.

## Acknowledgements

This research was supported by NIH R01 CA226285, LSU Collaborative Cancer Research Initiative, LSU Health Shreveport Office of Research, and LSU Health Shreveport FWCC Stimulus grants to X. Yu, LSU Health Shreveport FWCC Carroll Feist postdoctoral Fellowship to S. Cheng, and LSU Health Shreveport FWCC Carroll Feist predoctoral Fellowship to L. Li.

---

**sFig. 1.**
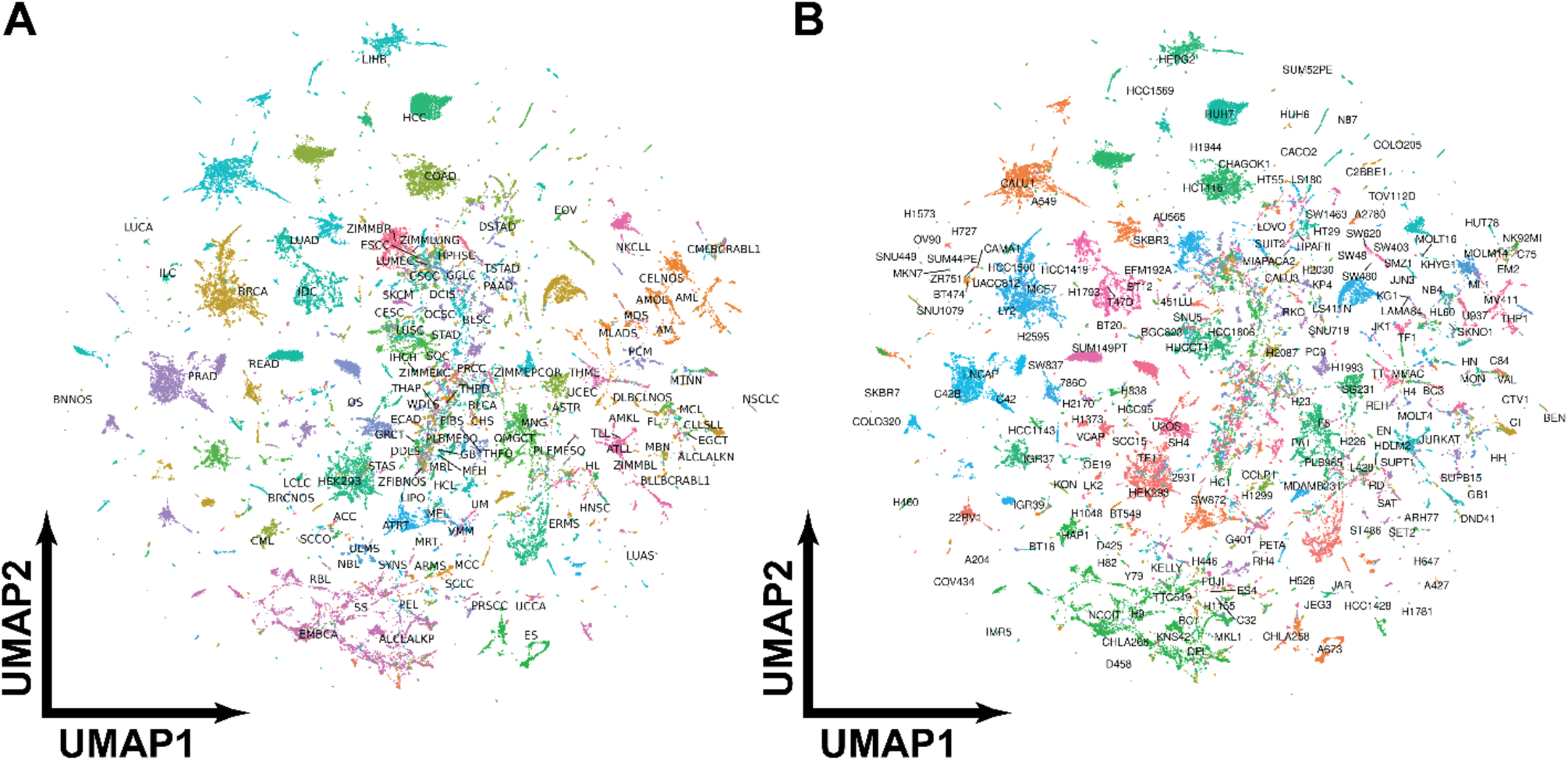
The UMAPs showed the samples had clustered location colored by their cancer types (A) and cell line names (B).

## Notes

### Competing Interest Statement

The authors have declared no competing interest.

